# Extracochlear Electric Stimulation – Toward Non-Invasive Hearing Restoration

**DOI:** 10.64898/2026.08.10.743874

**Authors:** Robert A. Hart, Patrick Hinz, Waldo Nogueira

## Abstract

**Background:** Hearing aids and cochlear implants (CIs) are the primary interventions for sensorineural hearing loss, restoring auditory function through amplification and intracochlear electrical stimulation, respectively. For those with residual low-frequency hearing, the combined electric-acoustic stimulation (EAS) has demonstrated superior speech perception, particularly in noisy environments, compared to either modality. However, CI surgery carries inherent risks, including postoperative hearing loss, which undermines EAS benefits and limits future rehabilitation options. To overcome these limitations, we propose a non-invasive alternative: extracochlear electric and acoustic stimulation (EEAS), delivering electrical stimulation via transcutaneous electrodes without surgery. Here, we present a first systematic investigation of non-invasive extracochlear electrical stimulation using ear canal electrode montages, evaluating its feasibility, perceptual effects, and key parameters across diverse hearing statuses.

**Methods:** We conducted a controlled, within-subject study with 15 participants: 5 with normal hearing (NH), 5 with high-frequency hearing loss (HI), and 5 with severe-to-profound deafness (PL). We used charge-balanced sinusoidal stimuli (125-4000 Hz) applied via an ear canal electrode and four return electrode montages, including contralateral ear canal, contralateral mastoid, ipsilateral mastoid, and forehead electrodes. Participants rated auditory sensations, including loudness, sound quality, and lateralization, as well as side effects on separate 0-10 scales, with current intensity increased up to 2 mA/cm². Thresholds and perceptual responses were analyzed across frequencies, electrode configurations, and hearing groups.

**Results:** Reliable auditory percepts were elicited across all groups. NH participants reported pure-tone sensations, whereas HI and PL participants perceived broadband, noise-like sounds. Loudness decreased with increasing frequency, particularly for HI and PL, with minimal responses in the high-frequency range. The current threshold increased with stimulation frequency, whereas the threshold expressed as charge per phase remained constant, suggesting that charge per phase primarily determines neural activation, whereas current amplitude is more closely associated with the intensity of auditory and side effect perception. Contralateral montages produced significantly higher loudness ratings than ipsilateral or forehead configurations. The forehead montage was poorly tolerated, leading to early termination due to discomforting side effects. Sound lateralization was predominantly central or bilateral with contralateral setups, while ipsilateral and forehead configurations yielded ipsilateral perceptions.

**Conclusions:** Non-invasive extracochlear electrical stimulation via ear canal electrodes is feasible and perceptually effective across a spectrum of hearing statuses. Perceptive outcomes are strongly influenced by electrode montage and residual hearing, with evidence of electrophonic excitation in NH individuals and electroneural activation in HI and PL participants. Contralateral mastoid electrode configurations offer the optimal balance of perceptual strength, tolerability, and spatial localization. These findings establish a critical foundation for the development of EEAS devices, demonstrating that non-invasive electrical stimulation can generate meaningful auditory percepts, paving the way for safe, accessible, and integrated hearing rehabilitation solutions. This work informs future EEAS developments and advances the path toward clinically viable, non-invasive cochlear stimulation.

## 2 INTRODUCTION

Hearing loss is among the most prevalent chronic health conditions worldwide and is associated with substantial social, cognitive, and economic consequences (Bainbridge and Wallhagen, 2014; Bisogno et al., 2021; Shukla et al., 2020; Wei et al., 2024). Although hearing aids remain the first-line treatment for most individuals with hearing loss, they do not fully restore auditory function and provide limited benefit for some patients, particularly those with severe high-frequency hearing loss or poor speech recognition(Marcos-Alonso et al., 2023; Moore, 2001). To address this unmet need while avoiding the risks associated with cochlear implantation, we propose a novel rehabilitation strategy: non-invasive extracochlear electric and acoustic stimulation (EEAS). EEAS combines electrical and acoustic stimulation without intracochlear electrode insertion, aiming to retain the benefits of combined stimulation while minimizing surgical invasiveness and preserving residual hearing.

Cochlear implants (CIs) provide reliable auditory rehabilitation for individuals with severe-to-profound sensorineural hearing loss through electric stimulation. Clinical outcomes demonstrate that CI implantation significantly improves speech intelligibility and overall quality of life, with the majority of recipients achieving clinically meaningful benefits (Gaylor et al., 2013; Tang et al., 2024). For individuals retaining residual low-frequency hearing, the combined use of a CI and a hearing aid, known as electric-acoustic stimulation (EAS), has been shown to further enhance speech perception, particularly in noisy environments, with even minimal residual hearing yielding substantial advantages over either modality alone (Büchner et al., 2009; Kiefer et al., 2005; Lenarz et al., 2013; Turner et al., 2004). Despite advances in surgical techniques and electrode design that minimize trauma during implantation, postoperative loss of residual hearing remains a significant risk, with complete preservation not guaranteed across all patients (Gerbert et al., 2024; Kopelovich et al., 2014; Zanetti et al., 2015). Median postoperative threshold shifts of approximately 10 to 15 dB have been reported at initial fitting, indicating immediate deterioration in residual hearing (Jurawitz et al., 2014). Furthermore, long-term follow-up studies have shown that around 20 % of EAS users lose sufficient low-frequency hearing to maintain the usage of the acoustic component of their speech processor within the first year, and up to 35% discontinue use of it within a decade (Gantz et al., 2016; Krüger et al., 2021). This loss of residual hearing not only reduces the benefit associated with EAS but also limits future access to emerging rehabilitation strategies that may depend on intact cochlear mechanics and functional hair cells. Unlike conventional EAS, which requires surgical insertion of an intracochlear electrode array, EEAS delivers electrical and acoustic stimulation without any or only minimal surgical intervention. The goal is to harness the synergistic advantages of combined stimulation while eliminating the risks and invasiveness of cochlear implantation.

Previous studies have investigated the use of extracochlear electrical stimulation as a means of improving speech understanding. However, these approaches primarily aimed to provide hearing through electrical stimulation alone and did not take residual acoustic hearing into account (Fourcin et al., 1983; Franz et al., 1989). Most approaches involve invasive electrode placements, such as in the round window niche, on the promontory, or within surgically created bony cavities, limiting their practicality and clinical applicability. Although these methods demonstrate the feasibility of extracochlear electric stimulation, they still require invasive surgical procedures, limiting their suitability for a practical hearing rehabilitation device. The long-term objective of this work is to develop a rehabilitation approach that combines electrical and acoustic stimulation without the need for cochlear implantation while maximizing the non-invasiveness of the system. Ear canal electrodes represent a promising alternative, offering a non-surgical pathway to extracochlear stimulation. They enable direct delivery of electrical currents to the cochlea through the ear canal, potentially bypassing the need for surgery. As a first step toward developing a fully non-invasive EEAS device, this study investigates the feasibility and perceptual effects of extracochlear electrical stimulation delivered via ear canal electrodes, systematically evaluating different return electrode montages and stimulation parameters. By avoiding surgical procedures, these electrodes provide a practical and easily deployable research platform, allowing for rapid evaluation of stimulation strategies and accelerating the exploration and optimization of extracochlear electrical stimulation concepts before progressing to minimally invasive approaches.

Although several studies have investigated electrical stimulation of the ear canal, most have focused on tinnitus management rather than hearing rehabilitation. Consequently, outcomes related to auditory perception, loudness perception, and differences across degrees of hearing loss remain largely unexplored (Kuk et al., 1989; Mielczarek and Olszewski, 2014; Suh et al., 2022; Vater et al., 2024; Zeng et al., 2019a). Tran et al. (2019) examined the influence of different electrode configurations on the estimated current reaching the cochlea; however, stimulation was limited to 500-Hz pulse trains presented below individual detection thresholds, precluding the evaluation of perceptual outcomes. Zeng et al. (2019b) performed a comprehensive investigation of sensory responses to transcranial electrical stimulation across multiple stimulation frequencies and electrode montages, including the usage of ear canal electrodes. While the study provided valuable insight into the dependence of auditory perception on stimulation parameters, the participant cohort consisted almost exclusively of normal-hearing listeners (with only two CI users), limiting generalizability to hearing-impaired populations. Although the authors reported qualitative differences in auditory perception between normal-hearing participants and CI users, the limited number of hearing-impaired participants precluded a broader assessment of hearing-loss-dependent effects. Moreover, auditory percepts in CI users have only been demonstrated using an ear canal-to-ear canal electrode montage, leaving it unclear whether alternative stimulation configurations could evoke comparable or superior perceptual responses. Suh et al. (2022) investigated electric hearing and tinnitus suppression in individuals with normal hearing and mild-to-moderate or high-frequency hearing loss, comparing ear canal and tympanic membrane stimulation. The study characterized hearing thresholds and pitch perception elicited by electrical stimulation. Yet, their analysis did not include systematic evaluation of return electrode configurations or detailed characterization of perceptual attributes such as loudness across frequencies or differences between hearing loss subgroups. More recently, Reinema et al. (2025) examined auditory perception and side effects of in-ear transcranial alternating current stimulation, varying electrode montage, frequency, and DC offset. However, the study was limited to normal-hearing participants and compared only two electrode configurations, leaving critical questions about optimal stimulation strategies in hearing-impaired populations unanswered. Given these gaps, further experimental data are urgently needed to identify effective electrode configurations and stimulation parameters for non-invasive EEAS. This requires a deeper understanding of the underlying physiological mechanisms across different degrees of hearing loss. Two primary mechanisms have been proposed: electroneural stimulation, involving direct activation of auditory nerve fibers by electric fields, and electrophonic stimulation, which results from electrically induced mechanical vibrations in the cochlea, likely mediated by outer hair cell activity and subsequent traveling wave propagation along the basilar membrane (Kipping et al., 2020; Moxon, 1971; Nuttall and Ren, 1995; Sato et al., 2017). In normal-hearing cochleae, auditory responses comprise contributions from both electrophonic and electroneural mechanisms, whereas electrophonic responses are abolished following the loss of functional hair cells, demonstrating their dependence on an intact cochlear function (Lusted and Simmons, 1988; Sato et al., 2016). While these mechanisms are well-documented in animal models, evidence in humans, particularly those with residual low-frequency hearing and high-frequency sensorineural hearing loss, remains sparse.

Based on this physiological foundation, we hypothesize that listeners with residual hearing will exhibit a combination of electrophonic and electroneural responses, with the relative contribution of electrophonic stimulation diminishing as hearing loss progresses. In contrast, individuals with severe-to-profound hearing loss are expected to rely predominantly on electroneural activation. The primary objective of this study was to systematically investigate auditory perception elicited by non-invasive extracochlear electrical stimulation delivered via ear canal electrodes across different return electrode montages, stimulation frequencies, and degrees of hearing loss. Specifically, we assessed perceived loudness, sound quality, and stimulation-related side effects to identify configurations that provide effective, comfortable, and perceptually meaningful auditory stimulation. Beyond parameter optimization, this work aims to advance understanding of the potential and limitations of non-invasive extracochlear stimulation, laying the groundwork for future development of combined EEAS rehabilitation devices, including sound-coding strategies and electrode configurations.

The remainder of this paper is organized as follows. Section 3 describes the study design, participants, stimulation system, and experimental procedures. Section 4 presents the experimental results, including perceptual responses, loudness perception, and side effects across electrode configurations and stimulation parameters. Section 5 discusses the implications of these findings for non-invasive extracochlear stimulation and the future development of combined EEAS rehabilitation devices. Finally, section 6 summarizes the key conclusions and outlines directions for future research.

## 3 MATERIAL AND METHODS

### 3.1 Participants

A total of 15 participants, aged between 23 and 77 years, were recruited from the German Hearing Center at the Hannover Medical School (MHH) to take part in two listening test sessions, each lasting approximately three to four hours. The sample included five individuals with normal hearing (NH group), five with high-frequency hearing loss (HI group), and five with severe-to-profound hearing loss across all frequencies (PL group). At the beginning of the first session, pure-tone hearing thresholds were measured (Figure 1) in a dedicated, acoustically treated chamber designed for clinical audiometric testing. Audiograms were obtained using an audiometer (Audio-Ton, Hamburg, Germany) and HDA-300 headphones (Sennheiser electronic GmbH & Co. KG, Wedemark, Germany). Participants with high-frequency hearing loss were typically bimodal users who had a CI on the non-target ear, with one individual using bilateral hearing aids. Among the participants with severe-to-profound hearing loss, one had a CI on the target ear and a normal hearing contralateral ear, while the remaining four were bilaterally implanted. Demographic details, including the duration and etiology of hearing loss, are summarized in Table 1. One participant (HI04) withdrew from the study after the first session due to unforeseen medical issues and was therefore unable to complete the second session. The study was conducted in accordance with the principles of the Declaration of Helsinki and received ethical approval from the Ethics Committee of the MHH. All participants provided written informed consent prior to participation.

**Figure 1.**
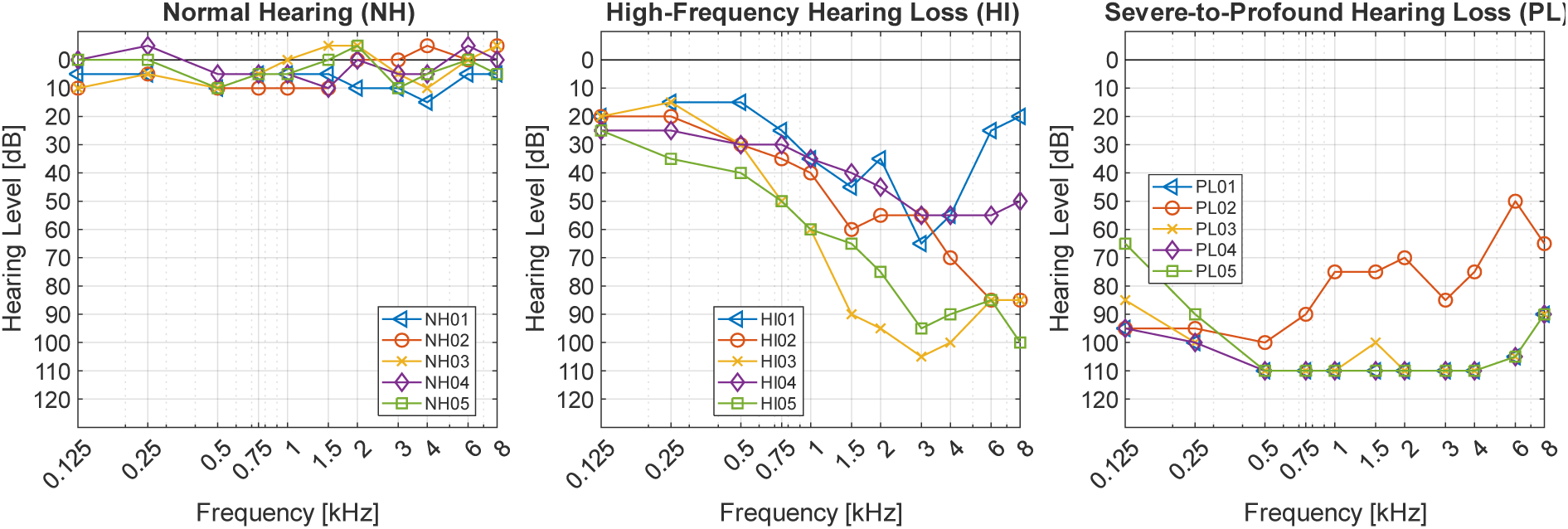
Pure-tone audiograms of the individual subjects for the three different hearing loss degree groups (Normal Hearing (NH); High-Frequency Hearing Loss (HI); Severe-to-Profound Hearing Loss (PL)).

**Table 1.** Demographics of the participants with three different degrees of hearing loss (Normal Hearing (NH); High-Frequency Hearing Loss (HI); Severe-to-Profound Hearing Loss (PL)).

| ID | Sex | Age | Stimulation Electrode Ear Side | Cause of Hearing Loss | Duration of Hearing Loss [years] | Implant |
| --- | --- | --- | --- | --- | --- | --- |
| NH01 | M | 36 | R | - | - | - |
| NH02 | M | 30 | R | - | - | - |
| NH03 | M | 27 | R | - | - | - |
| NH04 | M | 24 | R | - | - | - |
| NH05 | M | 35 | R | - | - | - |
| HI01 | F | 26 | L | Congenital, bilateral, Unknown | 26 | - |
| HI02 | F | 48 | L | Sudden sensorineural hearing loss (SSNHL) | 5 (ipsi), 26 (contra) | AB HiRes Ultra 3D HiFokus Mid-Scala |
| HI03 | M | 67 | L | Unknown | 14 | MED-EL Synchrony 2 Flex 28 S-Vector on contralateral side |
| HI04<sup>1</sup> | F | 63 | R | Sudden sensorineural hearing loss (SSNHL) on contralateral side | 16 (ipsi), 39 (contra) | Cochlear Nucleus CI622 on contralateral side |
| HI05 | M | 76 | L | Sudden sensorineural hearing loss (SSNHL) on contralateral side | 39 | MED-EL Synchrony 2 Flex 28 S-Vector on contralateral side |
| PL01 | M | 23 | R | Congenital, Unknown | 23 | Cochlear Nucleus CI24RE (CA) |
| PL02 | M | 32 | L | Unknown | 30 | Cochlear Nucleus CI522 |
| PL03 | M | 76 | R | Hereditary hearing loss | 51 | MED-EL Synchrony Standard |
| PL04<sup>2</sup> | M | 73 | L | Otosclerosis | 39 | Cochlear Nucleus CI24RE (CA) |
| PL05 | M | 77 | R | Unknown | 22 | Cochlear Nucleus CI522 |

<sup>1</sup> Subject had to stop after the first session due to medical reasons
<sup>2</sup> Subject had too small contralateral ear canal, preventing insertion of the tip trode

### 3.2 Experimental Setup

The experimental setup and procedures were adapted from those described by Zeng et al. (2019b) and are briefly outlined here. Transcranial alternating current stimulation was delivered between pairs of electrodes positioned at various locations on the subject’s head. Sinusoidal stimulation signals were generated using custom MATLAB scripts (MATLAB R2024b, The MathWorks, Inc., Massachusetts, USA) running on a Windows-based computer, enabling real-time adjustment of frequency and amplitude by the experimenters. The digital output was converted into an analog signal via a USB-connected data acquisition (DAQ) device (NI USB-6216, National Instruments Corp, Austin, USA). This analog signal was then used to drive a bipolar constant-current stimulator (Digitimer DS5, Digitimer Ltd, Welwyn Garden City, UK), which converted the voltage into a precisely controlled current waveform applied to the electrode pairs. During any phase of the experiment, stimulation could be immediately terminated by the experimenter through a manual control switch on the stimulator. To ensure safety, the peak current amplitude was limited to 2 mA/cm², in accordance with established safety guidelines. Prior to each testing session, the stimulation signals were verified using an oscilloscope and a 1 kΩ dummy load resistor connected to the stimulator output. This step confirmed signal integrity and prevented unintended current delivery. The electrodes used in the study consisted of gold foil tip trodes (ER3- 26A, Etymotic Research, Inc., Elk Grove Village, Illinois, USA), which were inserted into the ear canals of the participants. Additional electrodes were provided by tab electrodes (WhiteSensor 0315M EKG- Elektrode, Ambu, Copenhagen, Denmark), which were directly affixed to the skin at various locations on the skull. All electrode connections were secured using alligator clamps. A schematic representation of the complete experimental setup is shown in Figure 2.

**Figure 2.**
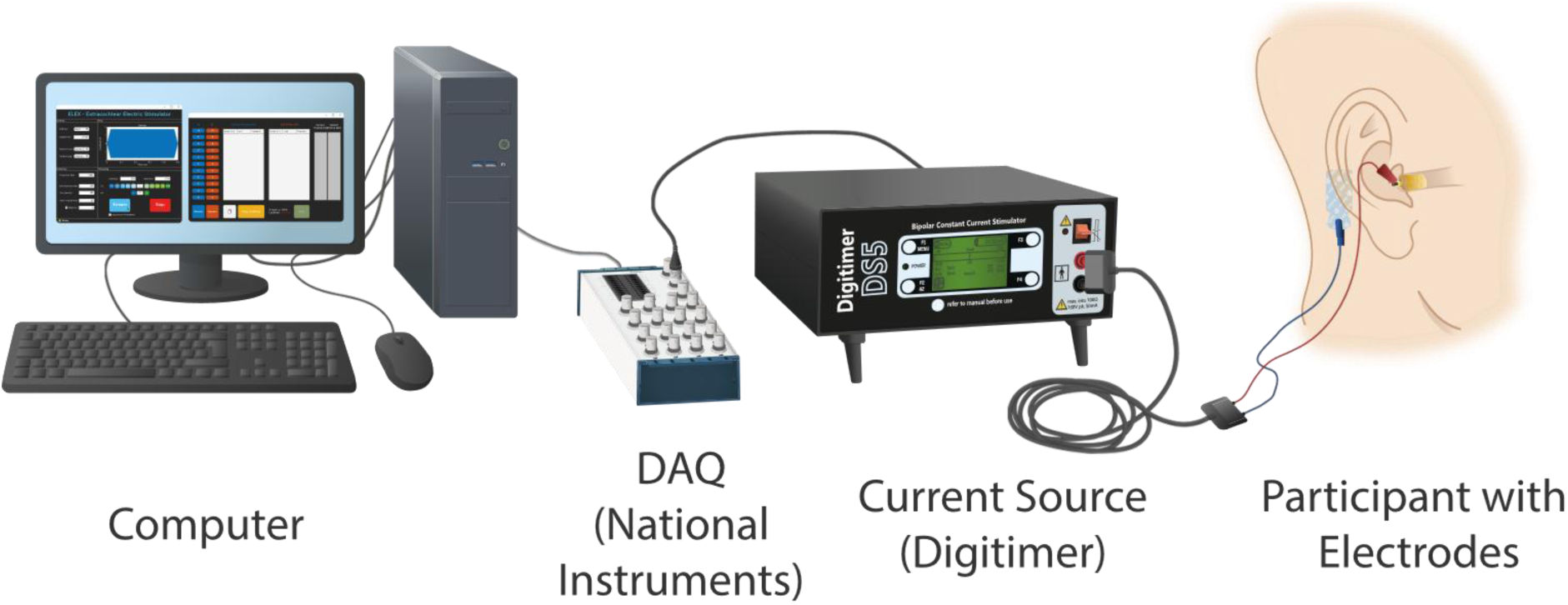
Schematic overview of the experimental setup using an ear canal tip trode and ipsilateral mastoid tab electrode. Signals are generated on a computer and transferred to a current source via a USB DAQ interface before stimulation is applied between electrode pairs.

### 3.3 Stimuli and Electrode Montages

Stimulation was delivered using 500 ms charge-balanced sinusoidal signals, each with 25 ms linear on- and 25 ms offset ramps to minimize transient effects. The stimulus frequencies tested were 125, 250, 500, 1000, 2000, and 4000 Hz. A total of four distinct electrode montages were evaluated, differing only in the placement of the return electrode. In all configurations, a stimulation electrode was placed in a single ear canal, serving as the target stimulation site for eliciting auditory sensations. The selection of a fixed ear canal electrode was guided by findings from Zeng et al. (2019b), who reported an increased auditory perception when using electrode montages that include ear canal electrodes compared to montages without. In participants with normal hearing, the ear with the better hearing thresholds was selected for stimulation. In individuals with hearing loss, the impaired ear was chosen as the stimulation site. The four return electrode positions investigated were: (a) the contralateral ear canal, (b) the contralateral mastoid, (c) the ipsilateral mastoid, and (d) a forehead-mounted tab electrode. These configurations are illustrated in Figure 3. Due to anatomical constraints, the contralateral ear canal electrode could not be placed in subject PL04, as the ear canal was too narrow, resulting in the exclusion of this condition for that participant.

**Figure 3.**
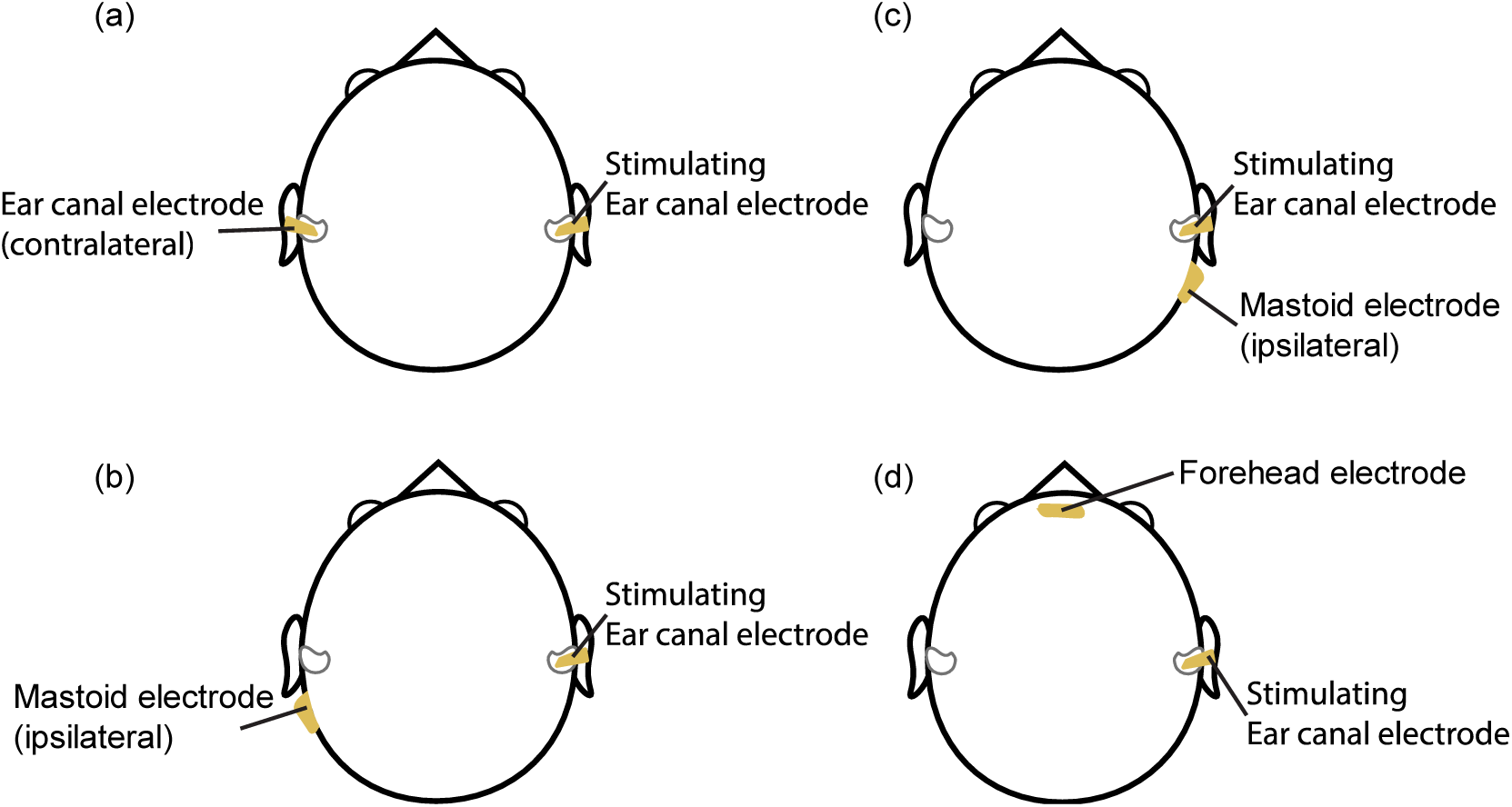
Electrode Montages, differing in return electrode location: (a) contralateral ear canal; (b) contralateral mastoid; (c) ipsilateral mastoid; (d) forehead.

### 3.4 Procedure and Loudness Scaling

All listening tests were conducted in a double-walled, soundproof booth with two examiners present. One was responsible for stimulus control and the other for recording responses. All electrodes were attached upfront. Before placing the ear canal electrodes, the external auditory canals were visually inspected for any obstructions. To ensure optimal conductivity, a conductive gel (GVB-geliMED GmbH, Bad Segeberg, Germany) was applied to the surface of the ear canal electrodes. For the tab electrodes, the corresponding skin areas were cleaned with alcohol wipes, and the electrodes were affixed directly using their integrated gel pads. Participants were seated comfortably in front of a computer screen, with the experimenter positioned facing them to monitor responses and maintain communication. Any CI processor or hearing aid was removed beforehand and was not worn during the test. Stimuli were delivered across a range of frequencies and electrode montages, beginning at 0 mA/cm² and increasing incrementally up to a maximum of 2 mA/cm² to ensure participant safety. Current intensity was increased in steps of 1 to 10 CU (current units), with progressively smaller step sizes at higher intensities to improve precision and reduce the risk of sudden discomfort. Equation 1 illustrates the relationship between stimulation current (µA) and current units (CU), based on the stimulation-level units commonly employed by major CI manufacturers. In CI devices, CU provides a logarithmic representation of current amplitude, enabling a wide dynamic range of electrical stimulation:

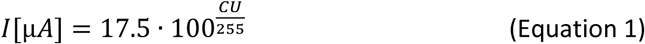

Participants were instructed to rate two distinct aspects after each stimulus: the perceived hearing sensation (HS) and any accompanying side effects (SE), each on a scale from 0 to 10. A score of 0 indicated no sensation or SE, while a score of 10 represented an unbearable or intolerable sensation. Responses included both the numerical intensity rating and a qualitative description of the perceived sensation, including the lateralization and characteristics. While side effects were reported according to their perceived location, sound sensations were classified as ipsilateral, contralateral or as central or bilateral, with respect to the stimulating ear canal side. The stimulation frequency and montage were maintained until either the sensation became too unpleasant or the maximum current of 2 mA/cm² was reached. The next electrode montage was then tested in a randomized order. To minimize order effects, the sequence of frequencies and montages was counterbalanced across participants. Prior to any stimulation, participants received a detailed explanation of the procedure and were given the opportunity to ask questions to ensure full understanding. To reduce the impact of sudden stimulation onset and prevent startle reactions, a 3-second countdown was displayed on the screen before each stimulus was delivered. To avoid expectancy or placebo effects, threshold detection was assessed using an ascending-descending procedure. When a sensation was first detected, the current was gradually reduced until the sensation disappeared. It was then increased again until the sensation reappeared. Although no formal pitch-matching procedure was conducted, side-by-side comparisons were performed using acoustic playback of sinusoidal tones at various frequencies when feasible.

## 4 RESULTS

### 4.1 Loudness Perception

Figure 4 displays the maximum HS loudness ratings reported by participants across different stimulus frequencies and electrode montages, stratified by hearing status. Within the NH group, maximum HS ratings remained relatively stable across frequencies and montages, with a median of 4 at 125 Hz and 250 Hz, followed by a gradual decline at higher frequencies. In contrast, the HI and PL groups exhibited a more pronounced frequency-dependent decline in maximum HS ratings. Sensation was most robust at the lowest stimulus frequency (125 Hz), with ratings decreasing sharply at higher frequencies. Notably, no perceptible hearing sensations were reported for the HI group at 2000 Hz, and only minimal responses were observed at 4000 Hz. The ipsilateral mastoid and forehead electrode montages showed the most extreme reductions in maximum ratings, particularly in the HI and PL groups, indicating a greater sensitivity to electrode placement in hearing-impaired individuals. Across all groups, 2000 Hz consistently yielded the lowest maximum ratings. The contralateral ear canal and contralateral mastoid montages were the only configurations that elicited detectable sensations in the HI group at frequencies above 1000 Hz, with the exception of 2000 Hz. In contrast, the ipsilateral mastoid and forehead montages failed to produce reliable responses in this group at higher frequencies. Given the small sample size (n = 5 per group), variability in the data was inherently high. However, the observed variability appeared greater in hearing-impaired participants, likely reflecting individual differences in residual cochlear function and current spread.

**Figure 4.**
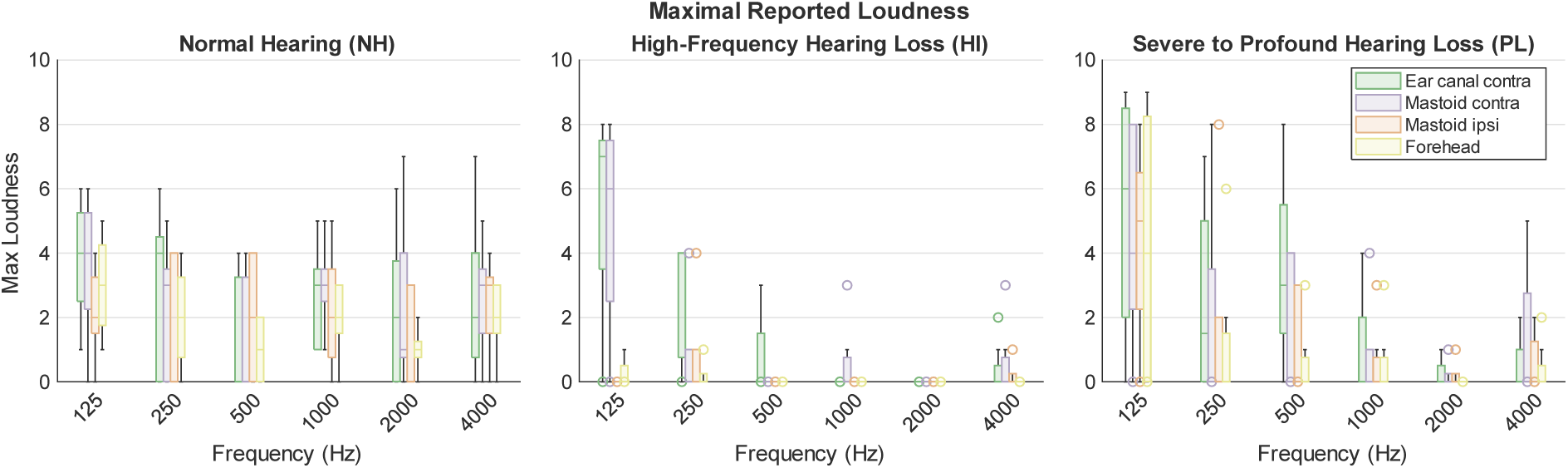
Maximum loudness or hearing sensations reported for different frequencies and electrode montages for individuals with Normal Hearing (NH), High-Frequency Hearing Loss (HI) and Severe-to-Profound Hearing Loss (PL). Horizontal lines within boxes show the median, box edges indicate 25^th^ and 75^th^ percentiles. Whiskers represent the data variability outside the interquartile range, while circles indicate outliers.

To assess the effect of electrode montage on maximum HS ratings, a Friedman test was conducted across the four montages. The analysis revealed a significant main effect of montage on perceived loudness (χ²(3) = 18.682, p < .001). Post-hoc pairwise comparisons were performed using Wilcoxon signed-rank tests, with p-values adjusted using the Holm-Bonferroni correction to control for multiple comparisons. Significant differences were found between the contralateral ear canal and both the ipsilateral mastoid (p = .039) and forehead (p = .020) montages. Similarly, the contralateral mastoid montage yielded significantly higher ratings than the ipsilateral mastoid (p = .018) and forehead (p = .020) configurations. In contrast, no significant differences were observed between the two contralateral montages (p = .698) or between the ipsilateral mastoid and forehead montages (p = .514). These findings are illustrated in Figure 5, which highlights the superior performance of contralateral electrode configurations in eliciting measurable hearing sensations, particularly in individuals with hearing loss. The results suggest that electrode placement plays a critical role in the effectiveness of extracochlear stimulation, with contralateral configurations offering a more reliable and robust perceptual response.

**Figure 5.**
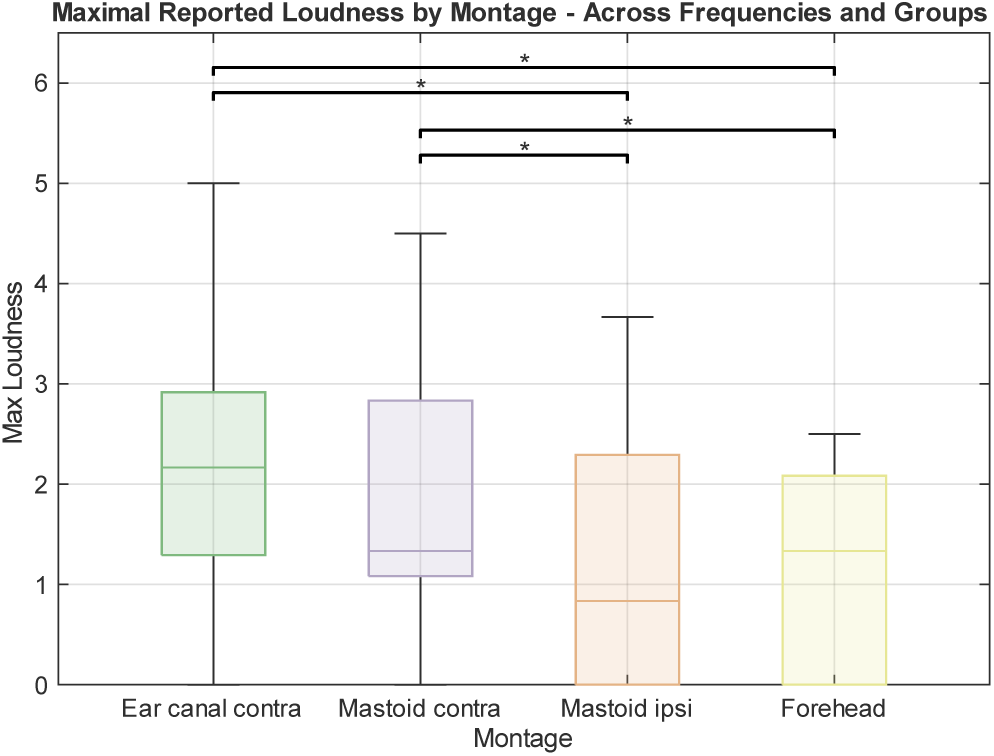
Maximum loudness or hearing sensations reported for different electrode montages across all participants. Horizontal lines within boxes show the median, box edges indicate 25^th^ and 75^th^ percentiles. Whiskers represent the data variability outside the interquartile range. Stars indicate statistically significant differences.

To analyze the relationship between stimulation current and perceived loudness ratings, individual rating profiles were interpolated to a uniform current grid with 0.01 mA resolution. Since ratings were recorded only when they changed during the experiment, ratings were assumed to remain constant between two consecutive measurements (zero-order hold interpolation). Thus, a reported rating was assigned and retained for subsequent current values until a change in rating occurred. Following interpolation, ratings were averaged across participants within each group at every 0.01 mA current step. If a participant was not measured beyond a certain current value, the corresponding data points were treated as missing values and excluded from the calculation of the group average at those current levels. The resulting group mean ratings are displayed as heatmap in Figure 6, where color encodes the average rating for each combination of stimulation frequency, current amplitude and electrode montage, with darker colors indicating higher perceived intensity ratings. Across all groups and montages, a consistent trend was observed: higher stimulus frequencies required greater current to elicit maximum hearing sensations. The frequency-dependent increase in required current was evident across all electrode configurations and participant groups. However, this trend was less pronounced in the HI group, where loudness declined sharply at frequencies above 500 Hz, as previously shown in Figure 4. An additional pattern emerged in the data: a dip in the maximum intensity rating at 250 Hz for most conditions, particularly in the NH group. At this frequency, the maximum perceived loudness was reached at lower current levels compared to other frequencies and often exceeded the ratings observed at 125 Hz and 1000 Hz. It is important to note that the shaded regions above the maximum reached current for each frequency in Figure 6 represent untested current levels, indicating that stimulation was not delivered beyond a certain current level because it produced intolerable side effects in any participant. As a result, the maximum current tested was limited by participants’ tolerance rather than by reaching a very loud sensation.

**Figure 6.**
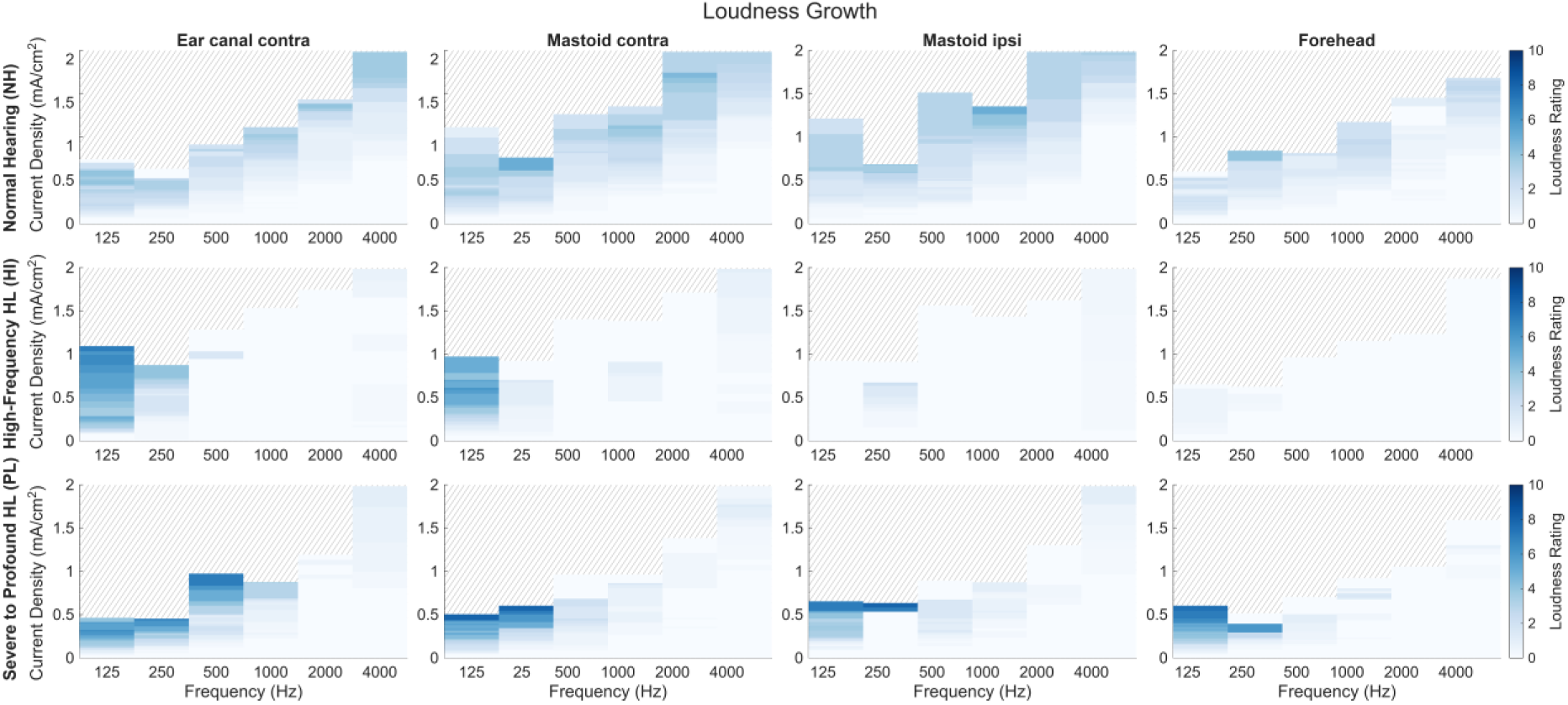
Heatmap showing the mean perceived loudness ratings as a function of stimulation frequency and current amplitude for each participant group (Normal Hearing (NH); High-Frequency Hearing Loss (HI); Severe-to-Profound Hearing Loss (PL)) and electrode montage. Individual ratings were interpolated to 0.01 mA current steps using a zero-order hold approach and averaged across participants at each current level. Higher loudness ratings are encoded with darker colors, lower ratings with lighter colors. Missing measurements were excluded from the averaging procedure. Shaded areas represent untested current levels.

### 4.2 Sound Lateralization

Figure 7 shows the percentage of sound lateralization across stimulus frequencies and averaged across frequencies for all three participant groups, focusing on the two contralateral electrode configurations. Lateralization was categorized as contralateral (perceived on the opposite side of the stimulating ear canal electrode), ipsilateral (perceived on the same side as the stimulating ear canal electrode), or central or both (a sensation perceived as originating inside the head or bilaterally from both ear sides). Across most frequencies, a high proportion of responses were classified as combined or central, particularly when using the contralateral ear canal montage. This condition also showed the highest percentage of contralateral-only lateralization, indicating that stimulation via the contralateral ear canal more consistently evoked sensations on the opposite side of the head. In contrast, the contralateral mastoid montage yielded a higher proportion of ipsilateral-only perceptions, especially between 500 Hz and 2000 Hz, where all participants reported sensations localized exclusively in the ipsilateral side. The HI group exhibited a distinct lateralization pattern with fewer ipsilateral-only responses and a predominance of combined or central perceptions. In contrast, the NH and PL groups showed more frequent ipsilateral-only localizations, particularly at lower frequencies. In this low frequency range, the proportion of contralateral-only perception was higher across all groups, while ipsilateral-only reports increased with higher frequencies, suggesting a potential frequency-dependent effect on sound lateralization. In addition to the categories shown in Figure 7, two instances of external perception were reported. Subject NH04 described a contralateral external HS at 1000 Hz when stimulated via the contralateral ear canal montage, perceiving the sound as originating outside the head, further away from the ear. Subject PL04 reported an ipsilateral external perception at 125 Hz using the contralateral mastoid montage, also describing the sound as originating beyond the head. These rare reports suggest that under certain conditions, extracochlear stimulation may evoke spatial perceptions that extend beyond the typical head-centered perception. The localization patterns for the ipsilateral mastoid and forehead montages were not included in Figure 7. The ipsilateral mastoid montage elicited only ipsilateral localizations across all frequencies and groups. The forehead montage produced exclusively ipsilateral sensations for all frequencies, except at 125 Hz in the HI group, where all participants reported central or both localizations, supporting the frequency-dependent effect observed in contralateral montages.

**Figure 7.**
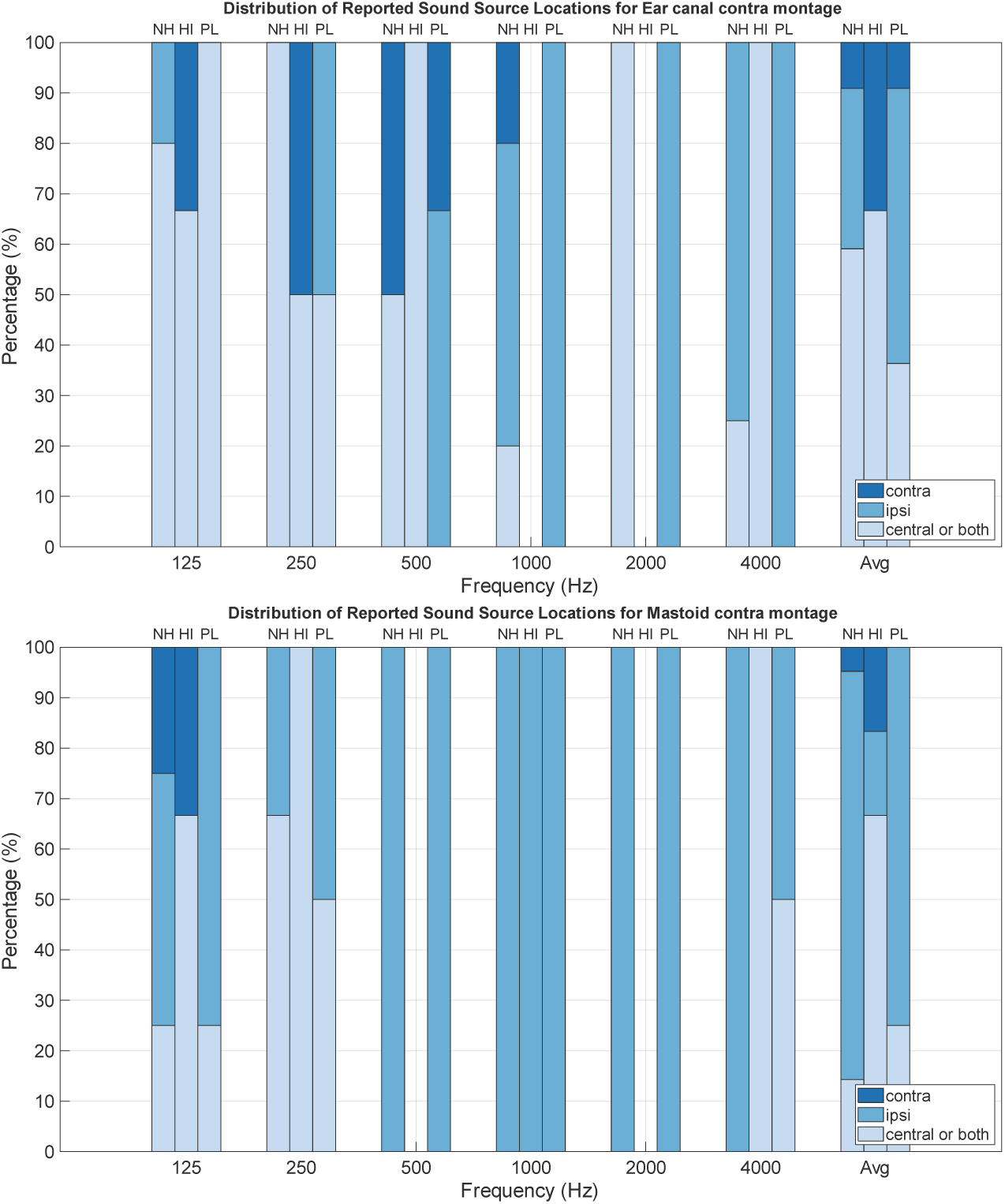
Percentage of lateralization for perceived hearing sensation (HS) elicited in Normal Hearing (NH), High-Frequency Hearing Loss (HI) and Severe-to-Profound Hearing Loss (PL) subjects for different electrode montages at different frequencies, as well as averaged across frequencies. Different shades of blue (dark to light) indicate percentages where the lateralization of hearing sensation was perceived only ‘ipsilateral’, ‘contralateral’ or on ‘both or central’ relative to the stimulating ear canal electrode.

### 4.3 Comments on Hearing Sensations

The auditory sensations elicited by extracochlear electrical stimulation in this study can be broadly categorized into two types: pure-tone or tonally clean perceptions, and broadband, noisy, or sound-like sensations. Table 2 summarizes the reported auditory sensations, categorized by participant group (NH, HI, PL), stimulus frequency, and electrode montage. Participants with NH predominantly reported pure-tone sensations that closely matched the frequency of the applied stimulus. In contrast, individuals of the HI and PL group more frequently described complex, broadband, or non-tonal sound qualities. Across all electrode montages, a general trend of increasing perceived frequency with higher stimulus frequency was observed. In the NH group, perceived frequency closely aligned with the actual stimulus frequency, indicating accurate frequency coding. However, the forehead electrode montage consistently yielded a perceptual shift, with perceived frequencies approximately doubling relative to the stimulus frequency, which has been previously reported in studies using non-invasive extracochlear stimulation (Suh et al., 2022; Zeng et al., 2019b). For the HI and PL groups, frequency perception was less precise. While some participants could distinguish low-, mid-, and high-frequency ranges, their descriptions were generally less specific. Instead of identifying exact tones, many described sensations using general terms such as “low,” “mid,” or “high” frequency, or compared the sounds to familiar environmental noises, such as car horns, doorbells, or buzzing sounds. Lower-frequency stimuli (125–500 Hz) were more frequently associated with recognizable sounds, whereas higher-frequency stimuli (1000–4000 Hz) often elicited vague or less distinct perceptions. Notably, the qualitative descriptions of auditory sensations did not differ significantly across electrode montages within each participant group. Despite variations in current distribution, the overall perceptual quality remained consistent across configurations, suggesting that the type of sensation (pure tone vs. noise-like) is more strongly influenced by hearing status than by electrode placement.

**Table 2.** Reported hearing sensations across frequencies for different electrode montages, divided into Normal Hearing (NH), High-Frequency Hearing Loss (HI), and Severe-to-Profound Hearing Loss (PL). Shaded areas represent no sound sensations.

| Group | Frequency (Hz) | Ear Canal Contra | Mastoid Contra | Mastoid Ipsi | Forehead |
| --- | --- | --- | --- | --- | --- |
| NH | 125 | low-frequency tone, hum, ~125Hz, sub-bass, pure tone | low-frequency tone, ~125Hz, pure tone | low-frequency tone, ~125Hz, amplitude modulated | low-frequency tone, hum |
|  | 250 | low-frequency tone, vibrating sound, ~200Hz, pure tone | low-frequency tone, hum, ~200Hz | low-frequency tone, high-frequency | low-frequency tone, low buzz, high-frequency tone, >250Hz |
|  | 500 | mid-frequency tone | mid-frequency tone | mid-frequency tone, high-frequency tone, ~500Hz, pure tone | mid-frequency tone, ~1kHz (octave shift) |
|  | 1000 | low-frequency tone, mid-frequency tone, ~2kHz (octave shift) | low-frequency tone, mid-frequency tone, ~2kHz (octave shift) | low-to mid-frequency tone, mid-frequency tone, ~2kHz (octave shift) | mid-frequency tone, ~2kHz (octave shift) |
|  | 2000 | mid-frequency tone, high-frequency tone, ~2kHz | mid- to high-frequency tone, high-frequency tone, ~2kHz | mid-frequency tone, high-frequency tone | low-frequency tone, mid-frequency tone, high-frequency tone |
| NH | 4000 | high-frequency tone | high-frequency tone, buzz sound | high-frequency tone | high-frequency tone, buzz sound |
| HI | 125 | low-frequency hum, doorbell ring, buzz | low-frequency, hum, doorbell ring, (bee) buzz, vibrating sound |  | vibrating sound |
|  | 250 | low- to mid-frequency hum, buzz, sound fork, noisy, vibrating sound | low-frequency sound, bell, high-pitched beep | noise | muffled tone |
|  | 500 | Buzz, vibrating sound |  |  |  |
|  | 1000 |  | (bee) buzz, vibrating sound |  |  |
|  | 2000 |  |  |  |  |
| HI | 4000 | high-frequency beep, bird-sound, vibrating sound | mid-frequency beep, vibrating sound | muffled tone |  |
| PL | 125 | low-frequency hum, high-frequency tone, buzz sound, cricket, alarm clock | low-frequency hum, high-frequency tone, buzz, dampened trumpet | low-frequency hum, high-frequency tone, buzz | low-frequency hum, high-frequency tone, buzz, dampened trumpet |
|  | 250 | low-frequency hum, speech-like feeling | low-frequency hum, speech-like feeling | low-frequency hum, vibrating sound | low-frequency hum |
|  | 500 | low-frequency hum, sound feeling, boom sound, buzz | low-frequency hum, muffled sound, muffled car honk | low-frequency sound, high-frequency sound, scream, noise, muffled church bell | low-frequency sound, car honk |
|  | 1000 | high-frequency sound | high-frequency sound | high-frequency sound | high-frequency sound |
|  | 2000 | high-frequency sound | high-frequency sound | mid-frequency sound |  |
| PL | 4000 | mid-frequency sound | high-frequency sound, hum, buzz | mid-frequency sound, hum, buzz, big/low car honk | high-frequency sound |

### 4.4 Side Effects

Participants were asked to report any SE experienced during stimulation and to rate their overall intensity. Figure 8 presents the percentage occurrence of SE across electrode montages and stimulus frequencies, aggregated across all participants. The most commonly reported SE were stinging and tingling sensations, primarily localized to the electrode sites, specifically the ear canal, mastoid, or forehead. These sensations were typically strongest at the return electrode location and were frequently cited as the most bothersome SE. Many participants compared the stinging sensation with needle stitches, pulling pain, or a feeling of electrical shock, while the tingling was often likened to a feeling of pressure or itching. Ear canal stinging was observed in all conditions, consistent with the fact that at least one ear canal electrode was used in every montage. Notably, the contralateral ear canal montage showed a higher incidence of ear canal stinging and tingling compared to other configurations, likely due to the involvement of two ear canal electrodes, thereby increasing the number of stimulation sites. Across frequencies, mastoid stinging decreased with increasing frequency, whereas forehead stinging increased, particularly at higher frequencies. The forehead electrode was reported by all participants as the most aversive site, with discomfort and pain often leading to early termination of stimulation. Muscle contractions were reported by many participants, particularly around the ear and facial regions. These contractions typically occurred toward the end of stimulation, just before the participant requested termination or the current limit was reached. They were more frequent at higher frequencies, except in the forehead montage, where such contractions were less commonly reported. Vibration sensations were also frequently reported, especially at lower frequencies, and were often localized to the return electrode site. Subject HI02 described vibrations that were either non-localizable or perceived as originating centrally within the head. Goosebumps were reported for multiple people, but often occurred only in a certain range of stimulation amplitude. In some subjects, the electrical stimulation could elicit a thermal sensation of coldness or warmth during the active stimulation. Notably, no participant reported dizziness, vertigo, or other symptoms suggestive of vestibular stimulation.

**Figure 8.**
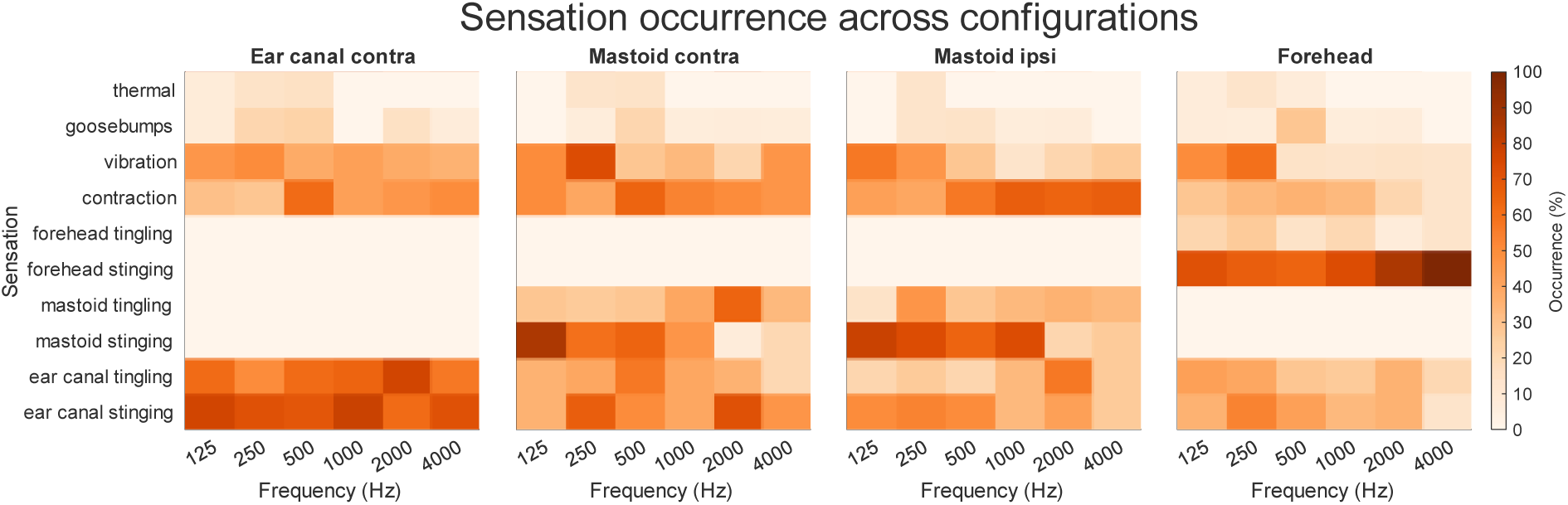
Percentage occurrence of different reported side effects at different frequencies and for different electrode montages, averaged across all subjects. Darker colors represent side effects with high occurrences, while lighter colors mean less probability of occurring.

Figure 9 illustrates the progression of SE intensity as a function of applied current, analogous to the loudness growth pattern shown in Figure 6. A similar trend is observed with higher stimulus frequencies requiring greater current amplitudes to reach the maximum tolerable SE level. The forehead montage consistently exhibited the lowest tolerance threshold, with SE reaching intolerable levels at relatively low currents, particularly at 500 Hz and above, and most prominently at 4 kHz. This condition was the first to trigger termination across multiple participants, confirming its high aversiveness. In contrast, the other montages allowed for higher current delivery before SE became unbearable. A notable pattern is the dip in current required to reach maximum tolerable SE at 250 Hz for many participants, mirroring the trend observed in loudness growth (Figure 6). At this frequency, participants reported that SE became intolerable at lower current levels compared to adjacent frequencies, suggesting a potential sensitivity peak in the 250 Hz range. Importantly, the PL group exhibited lower tolerance thresholds than both the NH and HI groups. Participants in the PL group reported higher SE ratings at lower current levels. As in Figure 6, the shaded regions above the maximum observed SE ratings represent untested current levels, indicating that stimulation was not provided beyond a certain threshold due to extreme discomfort or pain. Although higher current levels may enhance perceptual outcomes, such as increased loudness, they also elevate the risk of discomfort and pain, limiting the feasibility of further increases in stimulation intensity.

**Figure 9.**
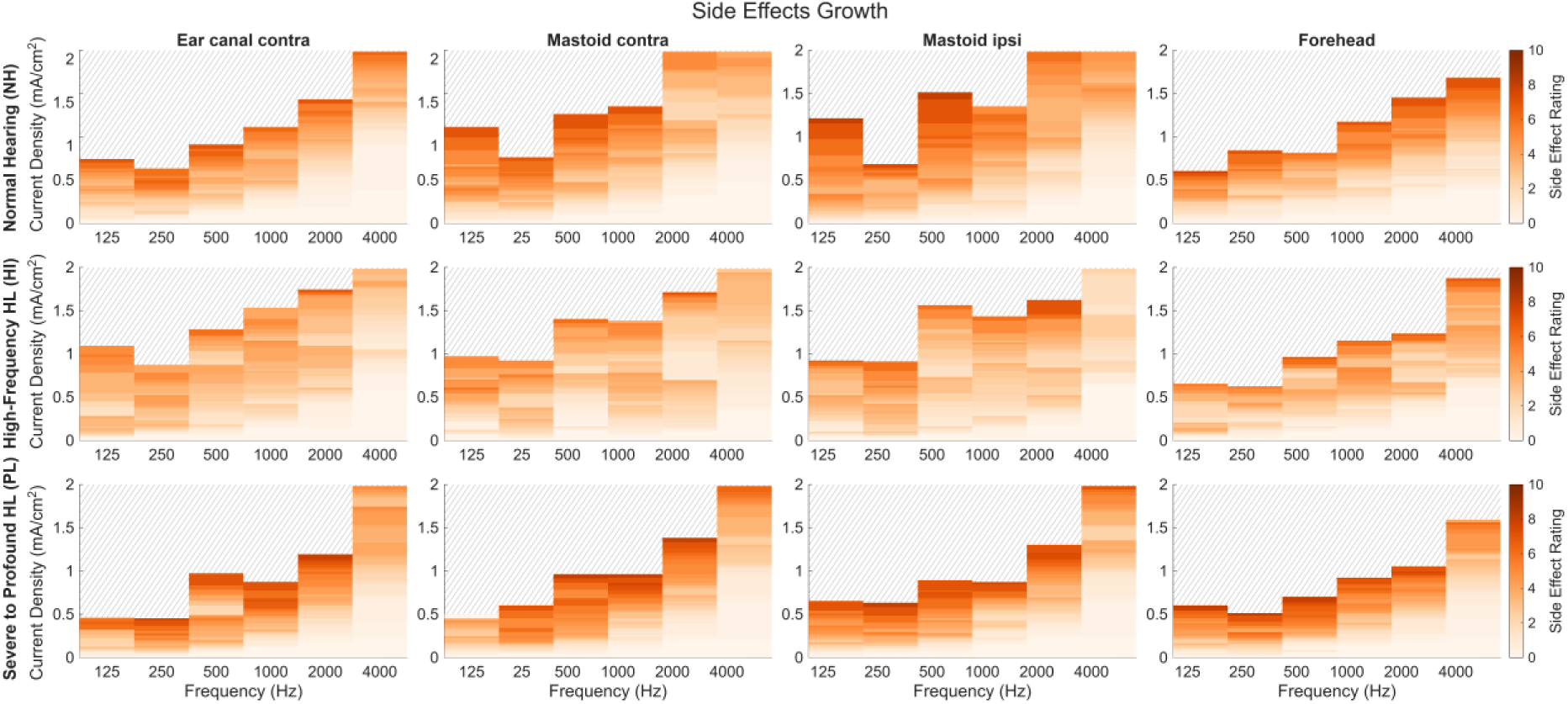
Heatmap showing the mean perceived side effect ratings as a function of stimulation frequency and current amplitude for each participant group (Normal Hearing (NH); High-Frequency Hearing Loss (HI); Severe-to-Profound Hearing Loss (PL)) and electrode montage. Individual ratings were interpolated to 0.01 mA current steps using a zero-order hold approach and averaged across participants at each current level. Higher side effect ratings are encoded with darker colors, lower ratings with lighter colors. Missing measurements were excluded from the averaging procedure. Shaded areas represent untested current levels.

### 4.5 Loudness and Side Effect Thresholds

Figure 10 presents the current necessary to elicit HS and SE thresholds across frequency and for different electrode montages (ear canal contra, mastoid contra, mastoid ipsi, forehead) and participant groups (NH, HI, PL). Across all configurations, HS thresholds occurred at higher current levels than SE thresholds. The growth of both thresholds followed an exponential trend with increasing stimulus frequency, and the gap between the two thresholds widened at higher frequencies. In the NH group, hearing thresholds closely followed the SE trend. In contrast, the PL group exhibited more variable threshold patterns, with local maxima appearing at certain frequencies depending on the electrode montage. Notably, in this group, the forehead montage yielded the lowest HS thresholds, which were often comparable to or even aligned with SE thresholds across most frequencies. The forehead montage again showed the lowest SE thresholds at 4 kHz, consistent with earlier findings. However, the overall SE threshold profile remained relatively similar across groups, indicating that the SEs were less dependent on hearing status.

**Figure 10.**
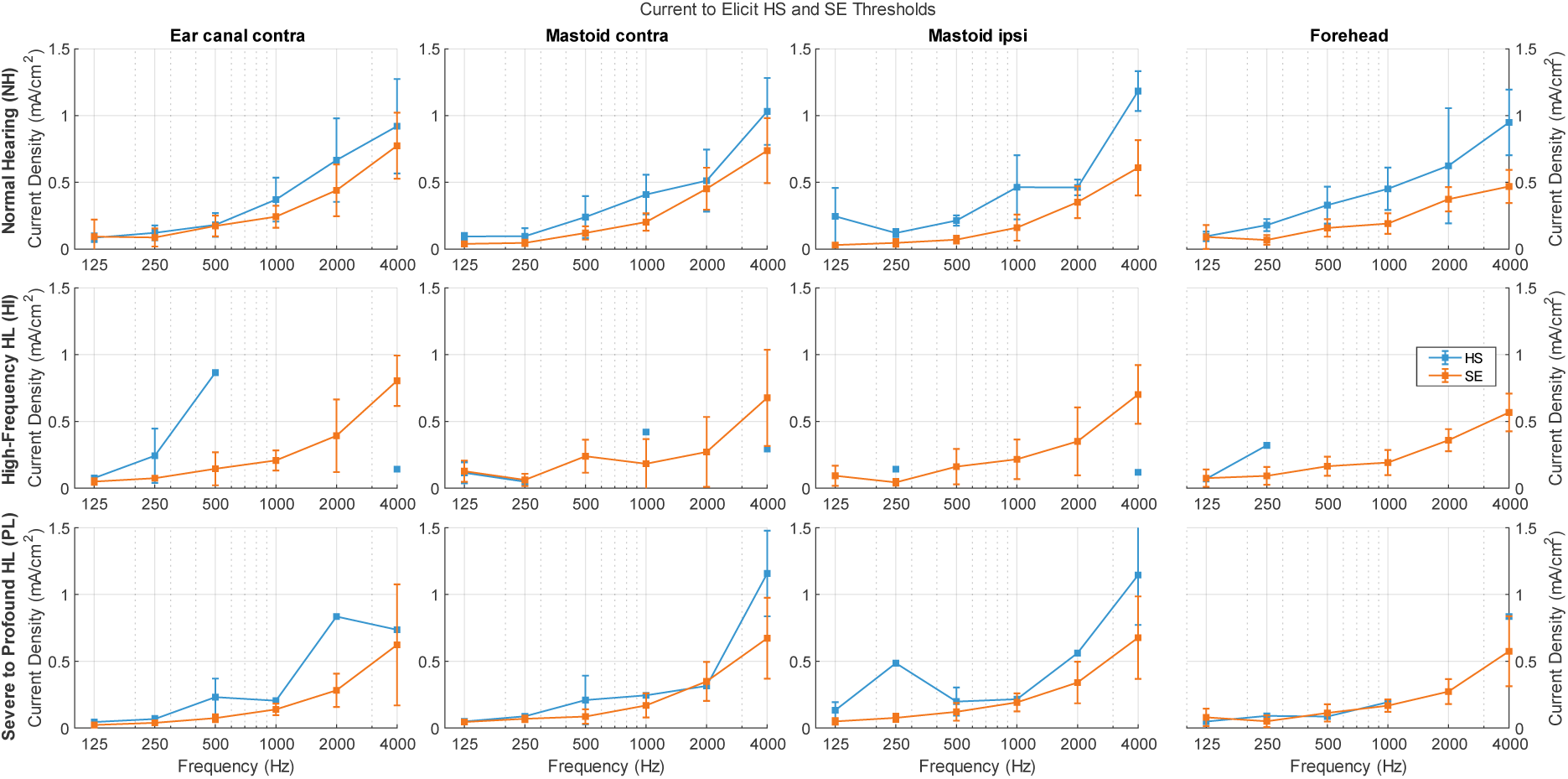
Average thresholds to elicit hearing (HS; blue) and side effects (SE; orange), for three groups of participants (Normal Hearing (NH); High-Frequency Hearing Loss (HI); Severe-to-Profound Hearing Loss (PL)). Squares represent group average and whiskers indicate ± standard deviation.

To further analyze the threshold behavior, the data were re-evaluated in terms of charge per phase, a key parameter in electrical stimulation that reflects the total charge delivered per half-cycle of a sinusoidal waveform. This is calculated using Equation 2:

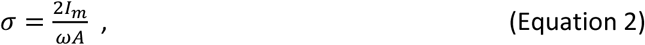

where *σ* is the charge density per phase, *I_m_* is the peak amplitude, *ω* the angular frequency and *A* is the electrode surface area. Figure 11 shows the charge density per phase required to reach thresholds of HS and SE. The average SE threshold across all measurements was at 152 nC/cm^2^ highlighted in the figure via the dashed horizontal line. The thresholds remained remarkably stable across frequencies and participant groups, suggesting a common physiological threshold for perceptual activation. This finding may help explain why higher frequencies are generally less effective in eliciting HS. To achieve the same charge per phase at higher frequencies, a greater peak current is required. At frequencies such as 4000 Hz, the necessary current often approaches or exceeds the safety limit of 2 mA/cm², making it difficult to reach the HS threshold.

**Figure 11.**
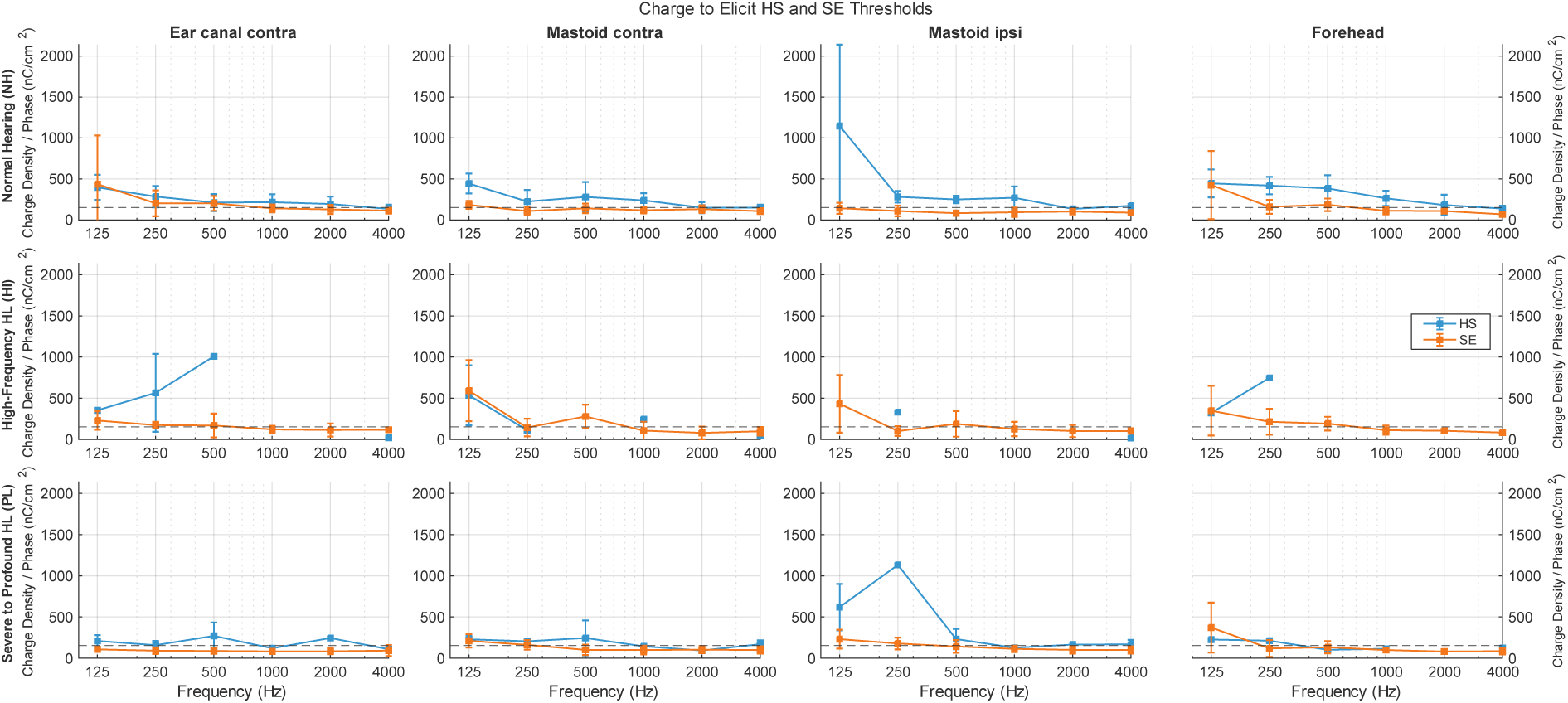
Average charge density per phase in nC/cm^2^ to obtain hearing thresholds (HS; blue) and side effect (SE; orange) thresholds, for three groups of participants (Normal Hearing (NH); High-Frequency Hearing Loss (HI); Severe-to-Profound Hearing Loss (PL)). Squares represent group average and whiskers indicate ± standard deviation. The dashed horizontal line represents the average SE threshold across all measurements.

## 5 DISCUSSION

This study investigated loudness perception and auditory sensation elicited by non-invasive extracochlear electrical stimulation across individuals with varying degrees of hearing loss. It extends prior research by systematically evaluating the effects of electrode montage and stimulation frequency while directly comparing perceptual responses among NH subjects and those with hearing impairment (HL and PL groups). The study was guided by two primary hypotheses. First, individuals from the NH group would predominantly exhibit electrophonic responses due to intact cochlear mechanics, whereas individuals from the PL group would rely primarily on electroneural activation and a combination of electrophonic and electroneural responses expected in the HI group. Second, contralateral electrode montages would yield louder sound sensations due to more favorable current pathways toward the cochlea. The results demonstrate the general feasibility of eliciting auditory sensations across all groups, but also indicate differences in perceptual characteristics and underlying response patterns depending on hearing status. Overall, the findings suggest that extracochlear electrical stimulation engages different mechanisms across populations and that electrode montage is a key factor shaping perceptual outcome.

### 5.1 Extracochlear electrical stimulation engages different auditory mechanisms across hearing status

The consistent loudness ratings and frequency-specific percepts observed in the NH group are consistent with electrophonic hearing. Participants in this group reported clear, tonal percepts that closely matched the applied stimulation frequencies, supporting the hypothesis that intact cochlear mechanics predominantly mediate auditory sensation during extracochlear electrical stimulation. In contrast, auditory percepts in the HI and PL groups differed markedly. Rather than reporting distinct tonal sensations, participants more commonly described broadband or noise-like sounds, suggesting that auditory perception in these groups was predominantly mediated by electroneural activation, even in individuals with residual low-frequency hearing. Our findings are in agreement with those of Zeng et al. (2019b), who reported sound perceptions of tonal quality and pitch matching to the stimulation frequency in normal-hearing and noise-like sounds in CI users, with pitch matching mainly to broadband noise or frequencies other than the stimulation frequency.

The effectiveness of extracochlear electrical stimulation also declined with increasing stimulation frequency in the HI and PL groups, as reflected by the low loudness ratings and the increasing number of participants who failed to perceive any auditory sensation (Figure 4). A similar trend of higher effectiveness at lower frequencies in normal-hearing using non-invasive extracochlear stimulation was also observed and has previously been reported, with lower current thresholds, particularly below 125 Hz (Reinema et al., 2025; Zeng et al., 2019b). However, the present study did not observe a comparable decline in perceived loudness across stimulation frequencies in the NH group compared to the HI and PL groups. The charge requirements for neural excitation may explain the reduced effectiveness at higher stimulation frequencies. Because charge per phase decreases with increasing stimulation frequency when current amplitude is held constant, higher frequencies require larger stimulation currents to achieve comparable neural activation. Consequently, perceptual thresholds expressed in current increased across all participant groups with increasing frequency. Studies in humans and cats have demonstrated a similar increase in auditory thresholds with shorter phase durations when using sinusoidal and biphasic pulses and a relatively constant charge per phase across frequencies required for eliciting hearing thresholds (Moon et al., 1993; Smith et al., 1995). In many cases, stimulation reached the predefined safety limit before higher loudness levels could be achieved, thereby restricting the maximum attainable loudness. Although higher current amplitudes might have increased the probability of eliciting auditory sensations, further increases were not justified because of the uncomfortable SE reported by the participants.

Although a similar trend of decreasing median loudness with increasing frequencies can be observed in the NH group, auditory perception remained more robust than in the HI and PL groups, with more frequent auditory sensations and consistently higher loudness ratings, even at higher stimulation frequencies. This difference is likely explained by the presence of intact cochlear mechanics. Electrophonic stimulation is thought to induce outer hair cell motility, generating mechanical vibrations that propagate as a travelling wave along the basilar membrane (Nuttall and Ren, 1995). As with acoustic stimulation, these vibrations can subsequently be amplified by functioning outer hair cells, thereby facilitating auditory perception (Oghalai, 2004). Consequently, NH participants may benefit from both electrophonic and electroneural mechanisms, whereas participants with hearing loss are likely to rely predominantly on electroneural activation because of reduced outer hair cell function. In principle, the residual low-frequency hearing observed in the HI group should still support electrophonic excitation in the apical region of the cochlea. However, the present findings indicate that its contribution was limited. Studies using intracochlear electrical stimulation have demonstrated that electrophonic thresholds increase with greater distance between the stimulation site and the region of electrophonic excitation (Sato et al., 2016). Given that the extracochlear electrode was positioned in the ear canal, it is plausible that the distance to the limited apical regions with preserved hearing was too large to evoke substantial electrophonic responses. This interpretation is further supported by the audiometric data (Figure 1), which show elevated low-frequency thresholds of approximately 20-30 dB in the HI group relative to the NH group, indicating that even the residual hearing region exhibited reduced cochlear sensitivity.

Current dispersion into surrounding tissues and bones may further contribute to the observed group differences. Back-telemetry measurements have shown that approximately 99% of the applied transcranial stimulation current is dissipated in surrounding tissues before reaching the cochlea (Zeng et al., 2019b). Because electrophonic stimulation is more effective than electroneural stimulation at relatively low current levels (Miller et al., 2006), only a limited portion of the residual cochlea may receive sufficient stimulation to generate electrophonic responses. This effect is likely to become increasingly important at higher stimulation frequencies, where larger currents are required to evoke auditory sensations but perceived SE and safety limits restrict the maximum deliverable current. Together, the reduced population of functioning outer hair cells, diminished sensitivity within the residual hearing region, and substantial tissue-induced current dispersion provide a plausible explanation for the differences observed between the NH and the two hearing-impaired groups.

A further indication of predominantly electroneural activation in the HI and PL groups is the nature of the reported auditory percepts. Participants of these groups generally described the stimulation as a complex, noise-like percept rather than a pure tone, as expected for electrophonic stimulation. However, perceptual differences between lower and higher stimulation frequencies suggest an effect of the temporal repetition rate at a constant stimulation site. This coarse distinction is consistent with the temporal rate coding of pitch during electrical stimulation, whereby perceived pitch varies as a function of stimulation frequency. Previous studies in CI users have shown that rate coding at more basal electrodes is typically associated with noisier percepts at low stimulation rates, with percepts becoming progressively cleaner as stimulation frequency increases (Landsberger et al., 2016). A comparable trend was observed in the PL group, where participants reported simpler auditory percepts at higher stimulation frequencies, consisting of fewer comparisons to more complex day-to-day sounds. Although this observation does not directly identify the site of neural activation, it is consistent with the possibility that the extracochlear stimulation predominantly excited more basal cochlear regions. If so, the limited contribution of more apical regions may further explain why participants in the HI group showed little evidence of electrophonic excitation despite retaining residual low-frequency hearing.

Moreover, the loudness growth functions (Figure 6) support the hypothesis that different activation mechanisms underlie auditory perception across groups. Compared to the NH group, both HI and PL groups showed more compressed loudness growth, consistent with a shift from predominantly electrophonic activation in normal hearing toward electroneural activation in people with hearing loss. This interpretation aligns with prior findings that electrophonic responses exhibit shallower amplitude growth than electroneural responses (Lusted and Simmons, 1988). While loudness growth alone cannot definitively distinguish between mechanisms, it provides complementary evidence for the proposed shift, confirming perceptual and frequency-dependent stimulation patterns.

Finally, the role of residual hearing or outer hair cell integrity in supporting electrophonic excitation remains unclear. Therefore, even mild auditory threshold elevations in the residual hearing of the HI group (Figure 1) may have reduced responsiveness to electrical stimulation, potentially limiting electrophonic contributions. Age differences between groups (NH: 30.4 years; HI: 56 years; PL: 56.2 years) may also have influenced outcomes, as age-related changes in neural responsiveness to electrical stimulation have been reported (Antonenko et al., 2021). However, the absence of clear electrophonic responses in the HI group cannot be interpreted as evidence that such responses cannot occur in this population. Electrophonic activation is highly dependent on stimulation configuration, electrode placement, and current spread (Lusted and Simmons, 1988). The current setup may have been suboptimal for eliciting such responses, and individualized parameter adjustments could yield different outcomes.

### 5.2 Inter-individual variability and group-dependent perceptual effects in hearing-impaired participants

A notable finding of this study was the lower overall perceptual outcome in the HI group compared to the PL group. This was unexpected, as the presence of residual hearing would typically be assumed to preserve, or at least not reduce, sensitivity to extracochlear electrical stimulation. However, the HI group exhibited the lowest average maximum loudness ratings and showed little to no perceptual responses at stimulation frequencies above 250 Hz.

The observed variability in perceptual responses may reflect inherent heterogeneity in extracochlear electrical stimulation, particularly in individuals with sloping hearing loss. The high within-group variability among HI participants observed in this study is consistent with prior reports of variable responses in similar populations, including no perception at higher frequencies (Zeng et al., 2019a). Factors such as individual anatomy, electrode placement, current spread, and age-related changes in neural responsiveness are known to influence electric field distribution and neural activation, contributing to variable perceptual outcomes (Antonenko et al., 2021; Laakso et al., 2015; Opitz et al., 2018). Adding to this, participant HI04 discontinued participation after the first session, limiting frequency coverage, while HI03 reached safety or discomfort thresholds early, restricting the dynamic range available for perceptual assessment. Such constraints likely further reduced the probability of observing reliable auditory sensations in these participants. Given that many sources of variability are intrinsic to the modality itself, the patterns observed in the current study align with expectations from prior work and suggest that the findings are representative of the broader challenges in extracochlear stimulation, rather than being uniquely attributable to sample size.

Differences in neural sensitivity and perceptual interpretation may further explain differences between the HI and PL group. Participants in the PL group consisted mainly of cochlear implant (CI) users who were experienced with electrically evoked auditory sensations and may therefore more readily interpret weak or ambiguous stimulation as auditory percepts. In contrast, participants in the HI group may still retain partially responsive auditory nerve fibers with spontaneous activity, which could reduce neural excitability due to refractory effects and thus reduce neural excitability due to refractory effects (Miller et al., 2006). This difference in perceptual learning and neural state may have contributed to the higher perceptual responsiveness observed in the PL group. The same group reported higher SE ratings at lower current levels, which might reflect altered neural excitability, heightened sensitivity to electrical inputs, or differences in peripheral and central processing in individuals with long-standing hearing loss and previous electrical stimulation knowledge. Subjective reports further support this interpretation. Several HI participants described difficulty distinguishing between vibrotactile SE and auditory sensations, with some noting that vibratory sensations occasionally transitioned into perceived sound as stimulation intensity. Future studies using objective measures such as cortical evoked responses could help clarify whether these percepts reflect true auditory nerve activation or differences in perceptual categorization. Whether perceptual sensitivity improves with training or repeated exposure could not be systematically examined in the present study and remains an open question.

Although electrode interaction effects from implanted CI arrays could in principle influence extracochlear current distribution (Tran et al., 2019), this is unlikely to explain the present findings given the similarity in stimulation conditions and the absence of systematic perceptual distortions specific to the PL group.

### 5.3 Electrode montage shapes auditory perception and sound lateralization

The present study demonstrates a clear effect of electrode montage on perceptual outcomes during extracochlear electrical stimulation. This finding is consistent with previous work showing that electrode configuration can critically influence the effectiveness of transcranial alternating current stimulation. For instance, altering only the position of the return electrode has been shown to significantly modify behavioral outcomes, with effects being most pronounced when the return electrode is placed contralateral to the primary stimulating electrode (Mehta et al., 2015). Similarly, intracochlear measurements have demonstrated that configurations placing one electrode near the target region and the return electrode on the contralateral side of the head maximize current delivery to the target site (Tran et al., 2019). In line with these findings, the present results indicate that contralateral montages generally produced stronger auditory percepts across stimulation frequencies and hearing groups.

Beyond perceptual strength, electrode montage also influenced the perceived spatial attributes of the elicited auditory sensations. Ipsilateral montages (mastoid and forehead) consistently produced lateralized percepts on the same side as the stimulation site. In contrast, contralateral configurations more frequently elicited bilateral, central, or contralaterally perceived sounds. This effect was more pronounced for the contralateral ear canal montage than for the contralateral mastoid montage and was observed across all participant groups. Frequency-dependent effects were also evident in the pattern of perceived laterality. In the contralateral ear canal configuration, mixed or bilateral percepts occurred across most tested frequencies, whereas the contralateral mastoid montage showed more restricted ranges of such effects, primarily at lower frequencies (≤500 Hz). Purely contralateral percepts were more common at low frequencies, occurring up to 1 kHz in the ear canal montage and only at 125 Hz in the mastoid montage. The high degree of spatial ambiguity in the ear canal configuration may reflect its more symmetrical current distribution, which could increase bilateral cochlear activation. In contrast, the asymmetry of the mastoid configuration may lead to more uneven current pathways and reduced bilateral spread.

### 5.4 Electrode montage limits auditory perception and stimulation tolerability via somatosensory side effects

Auditory perception in extracochlear electrical stimulation is fundamentally constrained by the maximum deliverable current, which is itself limited by stimulation-induced SE. In the present study, these SE consistently occurred at lower thresholds than auditory percepts, thereby heavily restricting the maximal achievable loudness. Electrode montage had a substantial impact on stimulation tolerability and SE profiles. Across configurations, participants reported non-auditory sensations including muscle contractions, vibrations, thermal sensations, and cutaneous tingling or stinging. However, the prevalence and intensity of these effects varied systematically with montage. In particular, tingling and stinging sensations at the return electrode site were frequently reported and often constituted the primary limiting factor for stimulation intensity. In many cases, these sensations occurred before auditory percepts reached higher loudness levels, thereby constraining the usable dynamic range. Among the tested configurations, the forehead montage was generally particularly poorly tolerated, with participants reaching discomfort thresholds at lower stimulation levels, making that montage unsuitable for practical use at higher stimulation frequencies and intensities. In contrast to findings by Reinema et al. (2025), no visual sensations such as phosphenes were reported under any condition or frequency. This is likely attributable to the selection of 125 Hz as the lowest stimulation frequency, which avoids the sub-100 Hz range shown to elicit phosphenes (Turi et al., 2013; Zeng et al., 2019b). Overall, these findings indicate that electrode montage not only shapes auditory percepts but also critically determines stimulation tolerability. Both factors should therefore be considered when optimizing electrode configurations for extracochlear stimulation paradigms. Increasing the surface area of the return electrode may reduce current density and improve comfort. Alternative stimulation configurations closer to the cochlea, such as tympanic membrane electrodes, may improve the balance of HS and SE and have been shown to elicit lower auditory thresholds and increase the likelihood of eliciting HS percepts (Suh et al., 2022). However, experimental setup remains challenging, as transcranial stimulation outcomes are highly sensitive to electrode placement, with even small positional deviations (≈1 cm) affecting reliability (Opitz et al., 2018).

### 5.5 Outlook and Future Work

Further studies are required to clarify the extent to which extracochlear stimulation interacts with cochlear hair cell function to induce electrophonic and electroneural responses. Such approaches may inform the development of future rehabilitation strategies. From this study’s outcome, an optimized processing aimed at electroneural activation seems reasonable, for example, through amplitude-modulated carriers and stimulation paradigms inspired by cochlear sound coding principles. In this context, biphasic pulsatile stimulation represents another promising avenue for further investigation. Follow-up studies should extend the present findings by investigating speech-relevant outcomes using combined EEAS, particularly in individuals with high-frequency hearing loss.

Electrode montage remains a critical design parameter for both perceptual efficacy and SE minimization. Depending on the intended application, different configurations may offer distinct advantages. Contralateral mastoid montages may provide a favorable balance between perceptual strength and spatial effects, whereas ear canal-to-ear canal configurations may be preferable when spatial localization is less critical. In contrast, forehead montages, which are frequently used in related stimulation studies and clinical applications such as tinnitus treatment, were associated with reduced tolerability and less favorable perceptual outcomes and should be used with more caution in other studies. These findings suggest that more optimized montage configurations may improve both efficacy and comfort in future extracochlear stimulation paradigms.

Last but not least, long-term effects of chronic extracochlear stimulation also remain to be systematically explored. Beyond the short-term perceptual effects, prolonged extracochlear stimulation may offer neuromodulatory benefits, such as supporting neuronal survival in the auditory pathway, which has been shown in animal models (Leake et al., 1995). In case training-related improvements in perceptual sensitivity are confirmed, the EEAS approach could additionally serve as a pre-implantation training tool, even for individuals with no residual hearing. Moreover, given its potential for tinnitus suppression (Vater et al., 2024), the device may have broader therapeutic applications beyond auditory rehabilitation alone.

## 6 CONCLUSIONS

This study investigated extracochlear electrical stimulation in 15 participants across three auditory profiles (normal hearing, high-frequency hearing loss, and severe-to-profound deafness) using multiple stimulation frequencies and electrode montages. The results demonstrate that extracochlear electrical stimulation can evoke auditory sensations across all groups, supporting its feasibility as a method for eliciting auditory percepts in both normal-hearing and hearing-impaired listeners.

Overall, perceptual outcomes were strongly dependent on stimulation frequency, with higher frequencies requiring increased current levels to elicit auditory sensations and showing reduced effectiveness, particularly in hearing-impaired participants. In addition, electrode montage significantly influenced both perceptual strength and spatial perceptual attributes, with contralateral configurations generally producing the highest loudness ratings but also increasing the likelihood of bilateral or spatially diffuse percepts.

The pattern of perceptual responses suggests different underlying mechanisms across auditory profiles. Responses in normal-hearing participants were most consistent with electrophonic activation, whereas hearing-impaired participants showed responses more consistent with electroneural stimulation under extracochlear ear canal stimulation. In addition, participants with high-frequency hearing loss exhibited slightly reduced perceptual outcomes compared to the severe-to-profound deaf group, while normal-hearing participants showed the most robust and consistent responses overall.

Taken together, these findings provide a baseline characterization of extracochlear electrically induced hearing across auditory profiles and stimulation configurations. They highlight the importance of stimulation frequency and electrode montage as key determinants of perceptual outcome, and they underscore the need for further optimization of stimulation parameters. These insights are relevant for the development of future auditory rehabilitation strategies combining extracochlear electrical and acoustic stimulation, as well as for improving the efficiency and tolerability of transcranial electrical stimulation paradigms.

## ACKNOWLEDGMENT

The authors gratefully acknowledge Benjamin Krüger from the Auditory Prosthetic Group, Hannover Medical School (MHH), for his valuable technical support. They also thank Constantino Dragicevic and Mechthild Meierott from the Institute of Neuroscience, Universitat Autònoma de Barcelona, Spain, for their assistance in setting up the experiments. Finally, the authors sincerely thank all study participants for their time, commitment, and valuable contribution to this research. This work is part of the READIHEAR project, which received funding from the European Research Council (ERC) under the European Union’s Horizon-ERC program (Grant agreement READIHEAR No. 101044753; PI: Waldo Nogueira). This work was also supported by Deutsche Forschungsgemeinschaft (DFG, German Research Foundation) cluster of excellence “Hearing4all” EXC 2177/1.

## Declaration of generative AI and AI-assisted technologies in the manuscript preparation process

The authors used ChatGPT (OpenAI) and a GWDG-hosted Qwen large language model (Alibaba Cloud) to improve the language, grammar, and readability of the manuscript. The authors reviewed and edited the content after using these tools and take full responsibility for the content of the published article.

## Notes

### Competing Interest Statement

The authors have declared no competing interest.

